# *Toxoplasma gondii Tg*MIF mediates the transmigration of extracellular parasites across the human placental barrier

**DOI:** 10.64898/2026.01.07.698173

**Authors:** Koen Brady Kleine, Guilherme De Souza, Lamba Omar Sangaré

## Abstract

The placenta is a critical biological barrier responsible for the healthy development of the fetus throughout pregnancy. However, the eukaryotic intracellular parasite *Toxoplasma gondii* (*T. gondii*) can cross the placental barrier by several mechanisms, including transmigration. Each year, approximately 190,000 infants are born worldwide with congenital infections caused by *T. gondii*, which can result in severe health complications or even death. Unfortunately, the molecular mechanisms underlying these tragic outcomes remain largely unclear, hindering the development of effective preventative strategies. To address this knowledge gap, we have developed a human *in vitro* placental barrier using trophoblast stem cells to study the transmigration of extracellular parasites across cellular tight junctions. Using this *in vitro* system, we found that *T. gondii* macrophage migration inhibitory factor (*Tg*MIF), a homolog of the human cytokine MIF, mediates extracellular parasite transmigration across the cellular tight junctions. Notably, *Tg*MIF, despite lacking a signal peptide, is actively excreted by the extracellular parasites. We found that the *Tg*MIF expression varies across *T. gondii* strains and positively correlates with the strain’s transmigration capacity. Mechanistically, *Tg*MIF distinctly induces the phosphorylation of extracellular signal-regulated kinase 1/2 and the dephosphorylation of focal adhesion kinase. These cellular modifications increase tight junction permeability, enhance parasite localization at these junctions, and facilitate subsequent transmigration. Furthermore, *Tg*MIF mediates transmigration independently of the host MIF receptor CD74, indicating the involvement of alternative receptors. Thus, our findings highlight *Tg*MIF as a critical effector that could mediate fetal infection in pregnant women via *T. gondii* transmigration across the placental barrier.

## INTRODUCTION

A pregnant woman can become infected through food or water contaminated with *T. gondii* infectious forms (cysts and oocysts) ^1^. Once inside the intestine, tachyzoites, the fast-replicative form, can rapidly disseminate to distant organs via the so-called Trojan horse mechanism ^2–6^, in which parasites use host immune cells as a vehicle. In the pregnant mouse model*, T. gondii* is detected in the spleen and mesenteric lymph nodes a few hours after infection, and in the maternal decidua 7 days post-infection. This suggests dissemination through the maternal blood ^7,8^. In humans, this blood route is also supported by cases of congenital infection, in which parasites are widely disseminated throughout all placental tissues ^9–13^. What is the structure of the human placental barrier?

The placenta has two interfaces that directly interact with the maternal decidua (anchoring villi) or with the maternal blood (floating villi) ^14,15^. The floating villi are made of migratory extravillous trophoblast (EVT), which anchors the placenta to the maternal decidua and remodels arteries to provide nutrient and oxygen-rich blood to the fetus ^16^. The syncytiotrophoblasts (STBs) form the floating villi; these multinucleated epithelial cells are polarized, expressing microvilli at their apical surfaces. STBs are bathed in maternal blood and have the largest surface area, which is critical for facilitating nutrient, gas, and waste exchange between mother and fetus ^14,15^. Beneath STBs reside the progenitor and mononucleated cytotrophoblast (CTBs) that differentiate into EVTs or STBs. Throughout the pregnancy, CTBs continuously replicate and fuse after the expression of the syncytialization marker syndecan-1 ^17,18^, replenishing the STB layer. Both STBs and CTBs constitute the placental barrier that protects the fetus from blood-borne pathogens ^16^. Indeed, STBs exhibit a striking resistance to *T. gondii* attachment to the plasma membrane and to intracellular replication ^19–22^. Human cases of congenital toxoplasmosis have revealed lesions in the floating villi; however, few or no parasites were observed in these lesions ^12,13^, confirming the resistance of this tissue to *T. gondii* intracellular replication. Therefore, how does *T. gondii* overcome the placental barrier?

There are over 156 distinct archetypal and non-archetypal strains of *T. gondii* ^23–26^ that can cause fetal infection. How frequently these strains effectively overcome the placental barrier is poorly understood and has been studied only to a limited extent. It is well known that extracellular parasites from several of these strains can actively transmigrate across the placental barrier and other biological barriers *in vitro* ^27,28^. In these studies, *in vitro* placental barriers were made from tumor-origin placental cell lines susceptible to pathogen infections, in which some parasites could cross after cell lysis ^29,30^. Transmigration requires the parasite’s active motility and the modulation of the host cell tight junctions ^31^. Within the archetypal groups found in North America, some strains exhibit differences in transmigration capacity, and this was attributed to a long-distance migratory phenotype (LDM) ^28^. During transmigration, *T. gondii* also exploits the host cell intercellular adhesion molecule 1 (ICAM-1) and induces the dephosphorylation of the focal adhesion kinase (FAK), which is essential for maintaining tight junction proteins such as ZO-1 ^31,32^. However, the parasitic effectors that mediate these cellular modulations or LDM are currently unknown.

The multifunctional cytokine macrophage migration inhibitory factor (*h*MIF) is highly secreted during the early stages of gestation, with levels gradually decreasing as pregnancy progresses ^33,34^. Notably, *h*MIF secretion during the initial phases of pregnancy appears crucial for protecting STBs against apoptosis ^35^. Interestingly, protozoan parasites, including *T. gondii*, *Plasmodium*, and *Entamoeba,* produce a functional homolog of the *h*MIF ^36^. Due to specific sequence differences, *T. gondii* MIF (*Tg*MIF) exhibits reduced tautomerase activity and lacks oxidoreductase activity compared to *h*MIF. Still, *Tg*MIF can stimulate IL-8 production in peripheral blood mononuclear cells (PBMCs), activate the ERK1/2 MAPK pathway in murine macrophages ^37^, and bind to *h*MIF receptor CD74 in an *in vitro* assay ^38^. Interestingly, *Tg*MIF functions remain unaffected by *h*MIF inhibitor ISO-1, highlighting the specificity of its structure and function ^37^. To date, no study has examined the capacity of *Tg*MIF to modulate cells at the placental barrier.

We used human trophoblast stem cells ^39^ to create a polarized, syncytialized placental barrier within a transwell system. This placental barrier enables us to focus exclusively on the ability of extracellular parasites to transmigrate, as they are resistant to intracellular replication. Our findings demonstrate that parasites deficient in *Tg*MIF exhibit significantly reduced transmigration capacity across cellular tight junctions. Furthermore, we demonstrate that the extracellular parasite actively excretes *Tg*MIF to modulate the ERK1/2 and FAK pathways, thereby facilitating the parasite’s localization at tight junctions and enhancing transmigration by increasing junctional permeability. Collectively, our research underscores the essential role of *Tg*MIF in modulating the human placental barrier during *T. gondii* transmigration.

## RESULTS

### Establishing an *in vitro* placental barrier made from human trophoblast stem cells

Tumor-origin cell lines have been previously used to generate *in vitro* placental barriers to study extracellular parasite transmigration ^31^. However, these cells cannot fully recapitulate placental cell resistance to *T. gondii* infection, such as attachment and intracellular replication restriction ^19,20^. We aimed to design a more accurate *in vitro* placental barrier system to study extracellular parasite transmigration. We grew human trophoblast stem cells (*h*TSCs) ^20,39^ under CTB conditions on a 12-well transwell membrane (0.9 cm^2^) with 8 µm pore size, coated with placental collagen IV + laminin 511, for 8 to 10 days. These cells had been used previously to develop an *in vitro* barrier on the basal side of a transwell system, differentiating into a mixed population of CTBs and STBs ^40^. As a control, non-barrier-forming human foreskin fibroblasts (HFFs) were also grown on transwells. After reaching complete confluence on the transwell system, we confirmed expression of the tight junction protein ZO-1 across the transwell culture (**Figure 1A**). We also confirmed the presence of syndecan-1, indicating that large areas of CTB have fused to form STB patches in the barrier (**Figure 1B**). The polarized and syncytialized CTB/STB, now called the placental barrier, presented a transepithelial electrical resistance (TEER) of 150-250 Ω*cm^2^, significantly higher than HFFs and similar to TEER values reported in prior literature using these cells ^40^ (**Figure 1C**). The placental barrier also exhibits less permeability to molecules such as 40 kDa FITC-Dextran (**Figure 1D**) than HFF. As expected, the placental barrier is more stringent to *T. gondii* intracellular growth (**Figure 1E**) and significantly limits parasite replication to one and two parasites per vacuole (**Figure 1F**) within 24 hours. To determine the transmigration capacity of extracellular parasites, we seeded 1×10^5^ wild-type (WT) RH-Luc parasites ^41^ onto the placental barrier. We counted the number that transmigrated to the bottom after 16 hours ^31^. Approximately 20% of the live parasites (viability determined by plaque assay) successfully transmigrated through the placental barrier, while more than 40% passed in HFF (**Figure 1G**). To examine whether changes in these barrier properties would affect the parasite transmigration, we stimulated the placental barrier with human cytokine interferon-gamma (IFN-γ) or human immunodeficiency virus (HIV) protein Nef-1, both of which alter TEER and permeability ^42–44^ (**Figures S1A and B**). Neither of these changes affected *T. gondii* transmigration (**Figure 1H**). Collectively, our *in vitro* system accurately mimics the placenta’s resistance to *T. gondii* intracellular replication, enabling us to assess only the extracellular parasite’s transmigration. Furthermore, it confirms that transmigration is not a passive process but is actively driven by parasitic effectors ^32^.

**Figure 1.**
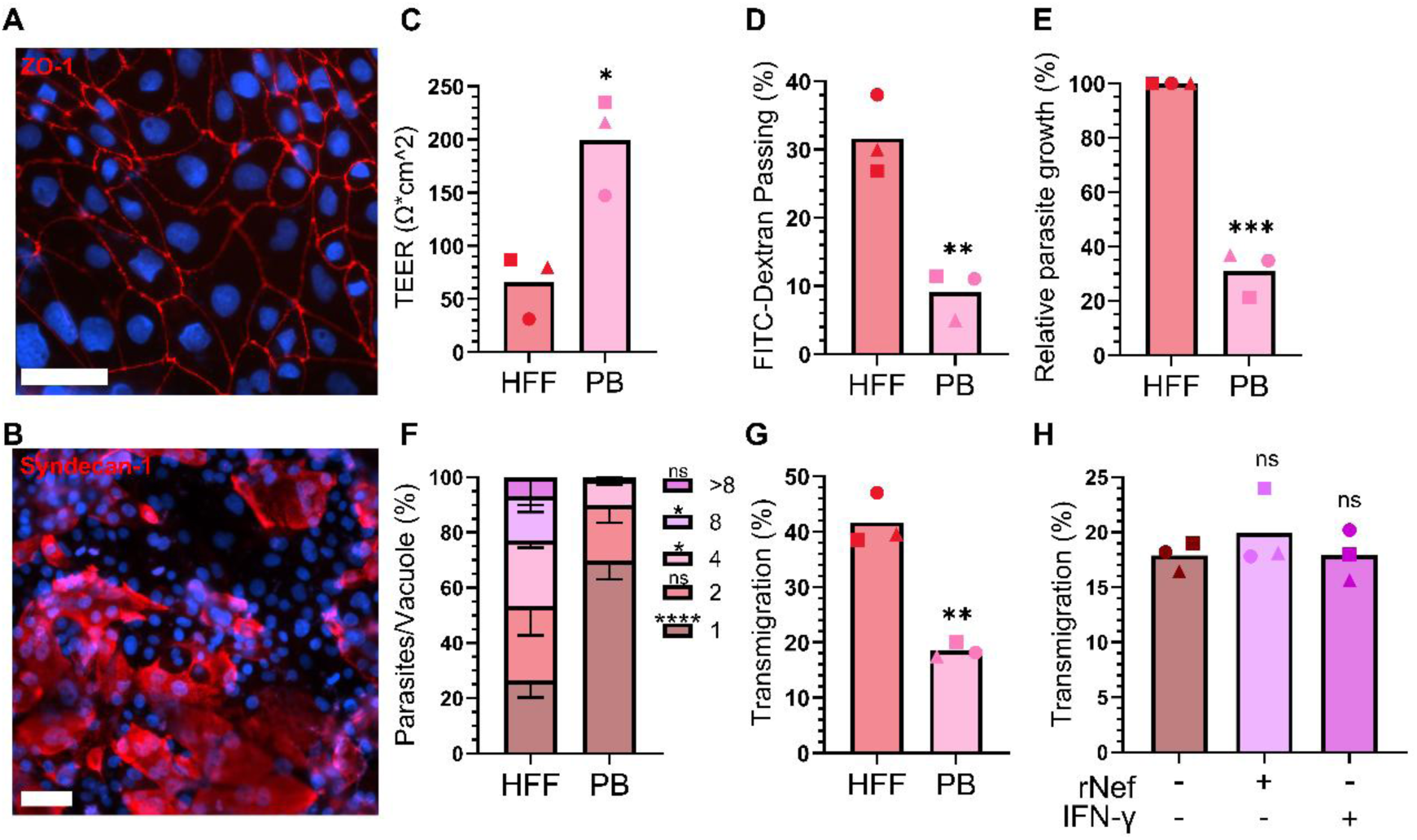
Validation of the human *in vitro* placental barrier (PB). **(A)** The tight junction structure in the placental barrier, stained with anti-ZO-1 antibody (red). Cell nucleus (blue). (Scale bar, 50 μm). **(B)** Syncytialization in the placental barrier, stained with anti-syndecan-1 antibody (red). Cell nucleus (blue). (Scale bar, 50 μm). **(C)** TEER measurements on HFF monolayer and the placental barrier. (*n=3*, means ± SD). Student’s t-test, \**P* = 0.0138. **(D)** Permeability of HFF monolayer and the placental barrier to 40kDa FITC-Dextran molecule. (*n=3*, means ± SD). Student’s t-test, \*\**P* = 0.0046. **(E)** RH-Luc parasite growth within 24 hours was quantified via luciferase assay in HFF monolayer and the placental barrier. (*n=3*, means ± SD). Student’s t-test, \*\*\**P* = 0.0001. **(F)** RH-Luc parasite growth within 24 hours was quantified by counting the number of parasites per vacuole in HFF monolayer and the placental barrier. (*n=3*, means ± SD). Two-way ANOVA with Sidak’s multiple comparisons, ns (not significant), \**P* < 0.05, \*\*\*\**P* < 0.0001. **(G)** RH-Luc parasite transmigration within 16 hours across HFF monolayer or the placental barrier. (*n=3*, means ± SD). Student’s t-test, \*\**P* = 0.0012. **(H)** RH-Luc parasite transmigration across the placental barrier, pre-treated or not with IFN-γ [100 ng/ml] or Nef [500 ng/ml]. (*n=3*, means ± SD). One-way ANOVA with Dunnett’s multiple comparisons, ns (not significant).

### *Tg*MIF mediates extracellular parasite transmigration across the human placental barrier

*T. gondii* expresses *Tg*MIF, a functional homolog of the cytokine *h*MIF, which is known to activate the ERK1/2 MAPK pathway ^37^. Extracellular parasite transmigration depends on the active modulation of host FAK and relies on ICAM-1 on the surface of the epithelial barrier ^31,32^. Interestingly, ERK1/2 MAPK, FAK, and ICAM-1 have interconnected functions that regulate various cell signaling pathways ^45^. Therefore, we hypothesize that *Tg*MIF mediates the transmigration of extracellular parasites. To explore this hypothesis, we generated an RH-Luc strain that is deficient in *Tg*MIF (Δ*Tg*MIF) using a CRISPR/Cas9 strategy ^5^. Compared to the WT, the Δ*Tg*MIF parasite shows no defects in replication using luciferase assay or parasite per vacuole count in HFF within 24 hours (**Figures 2A and B**) or in plaque formation after 5 days (**Figures 2C**). Furthermore, the absence of *Tg*MIF does not affect the parasite lethality in mice after intraperitoneal injection (**Figure 2D**). However, only 7% of viable Δ*Tg*MIF transmigrate across the placental barrier, which is significantly lower than WT parasites (**Figure 2D**). The transmigration capacity is restored to WT levels in Δ*Tg*MIF parasites that have been complemented with a copy of the *Tg*MIF gene inserted into the UPRT (uracil phosphoribosyltransferase) locus (**Figure 2E**), thereby confirming the role of *Tg*MIF in this process. The placental barrier expresses *h*MIF ^20,46,47^, which could have a synergistic effect alongside *Tg*MIF. To rule out this possibility, we first verified that Δ*Tg*MIF complementation with *h*MIF does not restore the transmigration capacity to WT parasite levels (**Figure 2E**). Then we used the specific inhibitor ISO-1 to inhibit *h*MIF excreted by the placental cells ^37^ and confirmed that it did not affect WT parasite transmigration (**Figure 2F**). Finally, we found that inhibiting the *h*MIF receptor CD74 with the neutralizing antibody Milatuzumab ^48^ did not affect WT parasite transmigration (**Figure 2G**). Therefore, *Tg*MIF is non-essential for the replicative cycle or virulence of *T. gondii* in mice; however, it mediates the transmigration of extracellular parasites across the human placental barrier, independently of *h*MIF and CD74 receptor.

**Figure 2.**
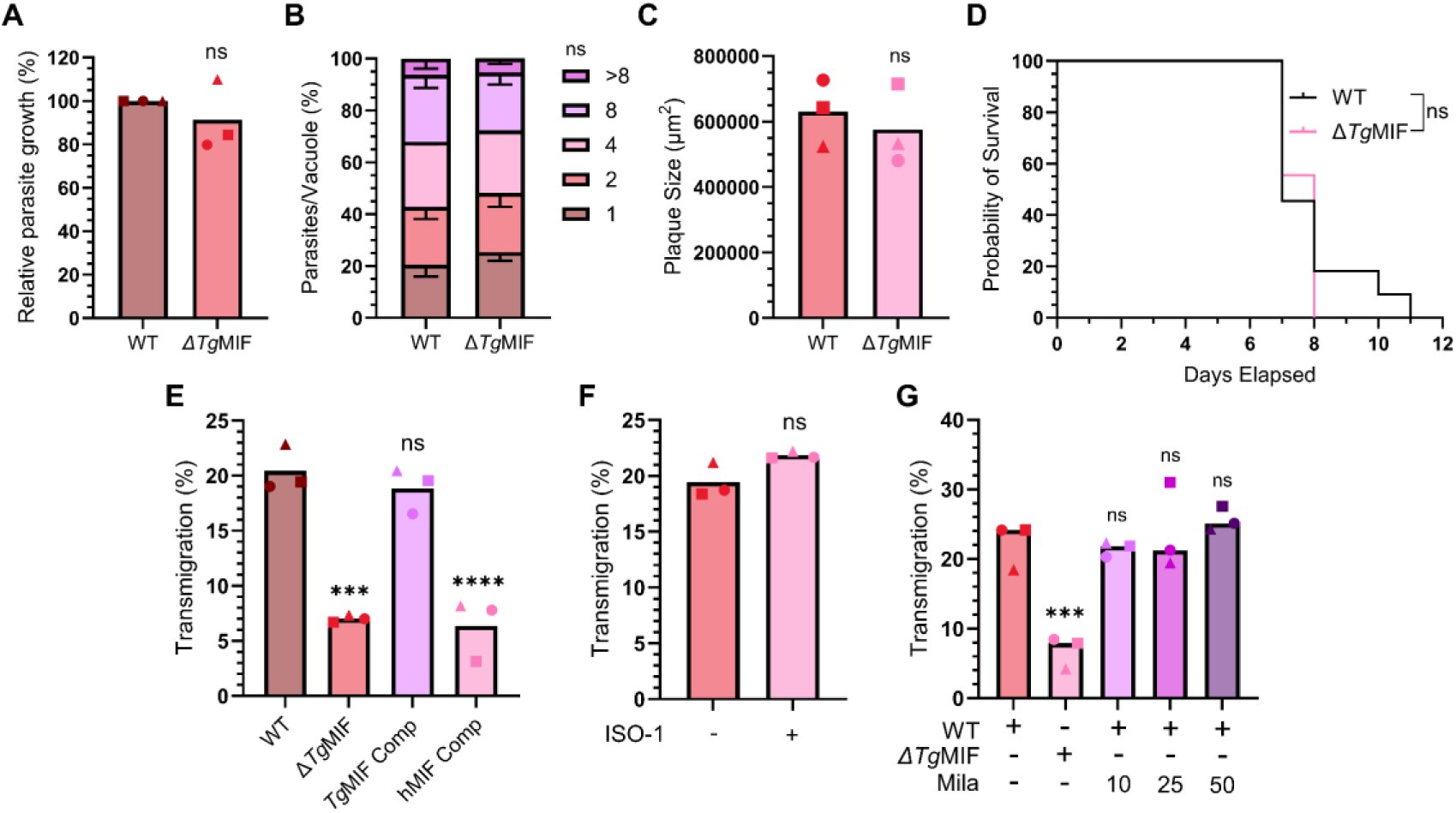
*Tg*MIF mediates the transmigration of extracellular parasites across the placental barrier. **(A)** WT (RH-Luc) and Δ*Tg*MIF parasite growth in HFF within 24 hours was quantified by luciferase assay. (*n=3*, means ± SD). Student’s t-test, ns (not significant). **(B)** WT and Δ*Tg*MIF parasite growth in HFF within 24 hours was quantified by counting the number of parasites per vacuole. (*n=3*, means ± SD). Two-way ANOVA with Sidak’s multiple comparisons, ns (not significant). **(C)** WT and Δ*Tg*MIF parasite plaque size was measured in the HFF monolayer after 5 days of infection. (*n=3*, means ± SD). Student’s t-test, ns (not significant). **(D)** Survival of CD-1 mice infected with 2,000 WT or Δ*Tg*MIF parasites. (*n=9*). Kaplan-Meier survival analysis, ns (not significant). **(E)** WT, Δ*Tg*MIF, *h*MIF compl, and *Tg*MIF compl parasites transmigration across the placental barrier. (*n=3*, means ± SD). One-way ANOVA with Dunnett’s multiple comparisons, ns (not significant), \*\*\**P* = 0.0001, \*\*\*\**P* < 0.0001. **(F)** WT parasite transmigration across the placental barrier treated or not for 30 minutes with ISO-1 [10 μM]. (*n=3*, means ± SD). Student’s t-test, ns (not significant). **(G)** WT and Δ*Tg*MIF parasites transmigration across the placental barrier pre-treated for 24 hours or not with the indicated μg/mL concentration of CD74 neutralizing antibody Milatuzumab (Mila). (*n=3*, means ± SD). One-way ANOVA with Dunnett’s multiple comparisons, ns (not significant), \*\*\**P* = 0.0009.

### *Tg*MIF distinctly modulates host ERK1/2 phosphorylation and FAK dephosphorylation

Recombinant *Tg*MIF has previously been shown to activate the ERK1/2 MAPK pathway and to induce IL-8 secretion in mouse bone marrow-derived macrophages and human PBMCs, respectively ^37^. To confirm that during infection, we infected the placental barrier with either WT parasites or with two distinct clonal isolates of Δ*Tg*MIF parasites for 45 minutes. We observed that the WT parasite strongly induces ERK1/2 phosphorylation, while both Δ*Tg*MIF clones fail to do so (**Figure 3A**). Therefore, we hypothesize that *Tg*MIF mediates extracellular parasite transmigration via the ERK1/2 MAPK pathway. To test this hypothesis, we used a chemical inhibitor of ERK1/2 phosphorylation, PD98059 ^49,50^, to treat the placental barrier. PD98059 treatment indeed decreases ERK1/2 phosphorylation even in the presence of WT parasites (**Figure S3A**). In the PD98059-treated barrier, we observed a significant decrease in WT parasite transmigration compared to the DMSO-treated and non-treated conditions, thereby confirming our hypothesis (**Figure 3B**). It is well established that *T. gondii* infection decreases FAK phosphorylation, and chemical inhibition of FAK phosphorylation with PF-573228 increased *T. gondii* transmigration ^32^. In fact, in our placental barrier treated with PF-573228, WT parasite transmigration is significantly higher than in the non-treated barrier (**Figure S3B**). Then, we hypothesize that *Tg*MIF mediates the dephosphorylation of FAK through the ERK1/2 MAPK pathway ^51^. We confirmed the first part of our hypothesis by treating the placental barrier with a recombinant *Tg*MIF protein (r*Tg*MIF) for 5 hours before western blot analysis. In a concentration-dependent manner, we observed that r*Tg*MIF significantly decreased FAK phosphorylation (**Figures S3C and D**). Following this, we treated the placental barriers with PD98059, DMSO, or left them untreated, and infected them with either WT or Δ*Tg*MIF parasites for 5 hours. Subsequently, we conducted Western blot analysis. Regardless of whether ERK1/2 MAP is inactivated, WT parasites significantly decreased FAK phosphorylation, while Δ*Tg*MIF parasites did not exhibit this function (**Figures 3C and D**). Consequently, *T. gondii* transmigration across the placental barrier relies on the modulation of ERK1/2 and FAK phosphorylation, both of which are distinctly mediated by *Tg*MIF.

**Figure 3.**
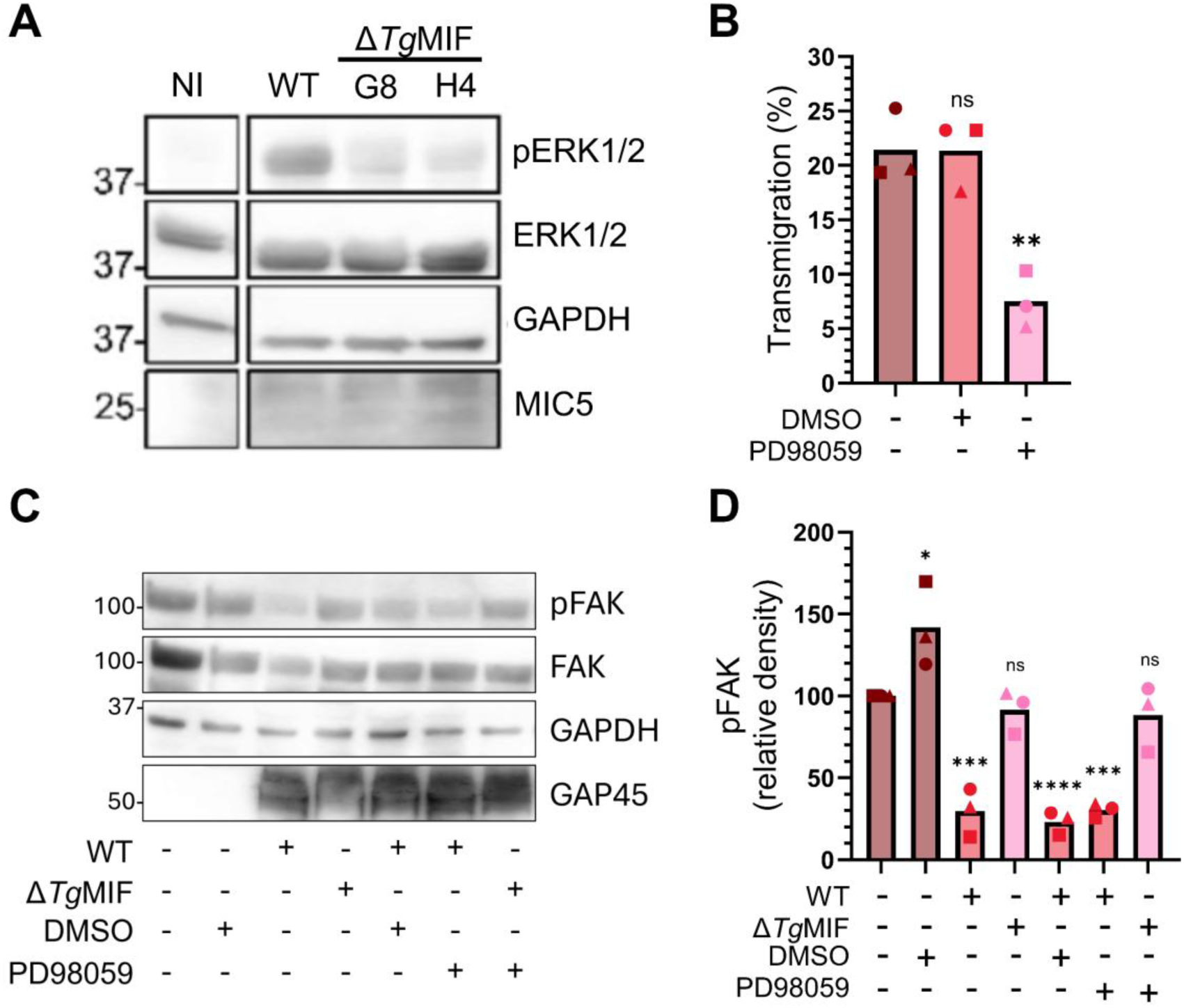
*Tg*MIF modulates the ERK MAPK pathway and induces FAK dephosphorylation. **(A)** SDS-PAGE analysis of lysates from placental barriers infected or not (NI) for 45 minutes with WT (RH-Luc) or Δ*Tg*MIF parasite, blotted with anti-phospho ERK1/2 (pERK1/2), anti-ERK1/2, anti-GAPDH (cell loading control), and anti-MIC5 (parasite loading control), as well as their respective secondary HRP-antibodies. **(B)** WT parasite transmigration across the placental barrier pre-treated 24 hours or not with PD98059 [50 μM] or vehicle DMSO. (*n=3*, means ± SD). One-way ANOVA with Dunnett’s multiple comparisons, ns (not significant), \*\**P* = 0.0026. **(C)** SDS-PAGE analysis of lysates from placental barriers infected pre-treated 24 hours or not with PD98059 [50 μM] or DMSO vehicle, and infected or not for 5 hours with WT or Δ*Tg*MIF parasite. The blotting was performed with anti-phospho FAK (pFAK), anti-FAK, anti-GAPDH (cell loading control), and anti-GAP45 (parasite loading control), along with their respective secondary HRP antibodies. **(D)** Relative quantification of pFAK in each condition from (C) after normalization with GAPDH. (*n=3*, means ± SD). One-way ANOVA with Dunnett’s multiple comparisons, ns (not significant), \*\*\**P* < 0.001, \*\*\*\**P* < 0.0001.

### *Tg*MIF mediates extracellular parasite localization at the cellular tight junction

*h*MIF enhances ICAM-1 expression in various cell types ^52,53^, and ICAM-1 plays a crucial role in the transmigration of *T. gondii* across polarized endothelial and epithelial barriers ^8^. We hypothesize that *Tg*MIF promotes upregulation of ICAM-1 expression at the placental barrier via the ERK1/2 MAPK pathway ^45,54^. We infected the placental barriers with either WT or Δ*Tg*MIF clones and performed Western blot analysis. Our results showed no difference in ICAM-1 expression between WT and Δ*Tg*MIF clones (**Figure 4A**). However, when the placental barrier was treated with an ICAM-1-specific neutralizing antibody ^31^, we also observed a significant decrease in WT parasite transmigration capacity (**Figure 4B**). To mediate leukocyte adhesion at the cellular tight junction, ICAM-1 is associated with other CAM proteins, such as VCAM-1 ^55,56^. We hypothesize that *Tg*MIF mediates extracellular adhesion and tight junction localization. To test this, we seeded WT or Δ*Tg*MIF parasites onto the placental barrier for 5 hours. The barrier was then fixed and stained with ZO-1 for immunofluorescence. We captured images of random fields and quantified the number of WT and Δ*Tg*MIF parasites positioned at the tight junction **(1)**; within 2 μm of a tight junction **(2)**; and at distances greater than 2 μm from a tight junction **(3)** (**Figure 4C**). Our data reveal that a significantly higher percentage of WT parasites (35.75%) localize at tight junctions compared to Δ*Tg*MIF parasites (22.57%). Conversely, a larger proportion of Δ*Tg*MIF parasites (38.05%) is found at a greater distance from the junctions than WT parasites (28.25%) (**Figure 4D**). Notably, when seeded on barriers stimulated with 100 ng/mL of r*Tg*MIF, the localization of Δ*Tg*MIF parasites at cellular tight junctions significantly increased (**Figure 4D**). Collectively, these findings highlight the critical roles of ICAM-1 and *Tg*MIF in facilitating extracellular parasite adhesion and localization to cellular junctions.

**Figure 4.**
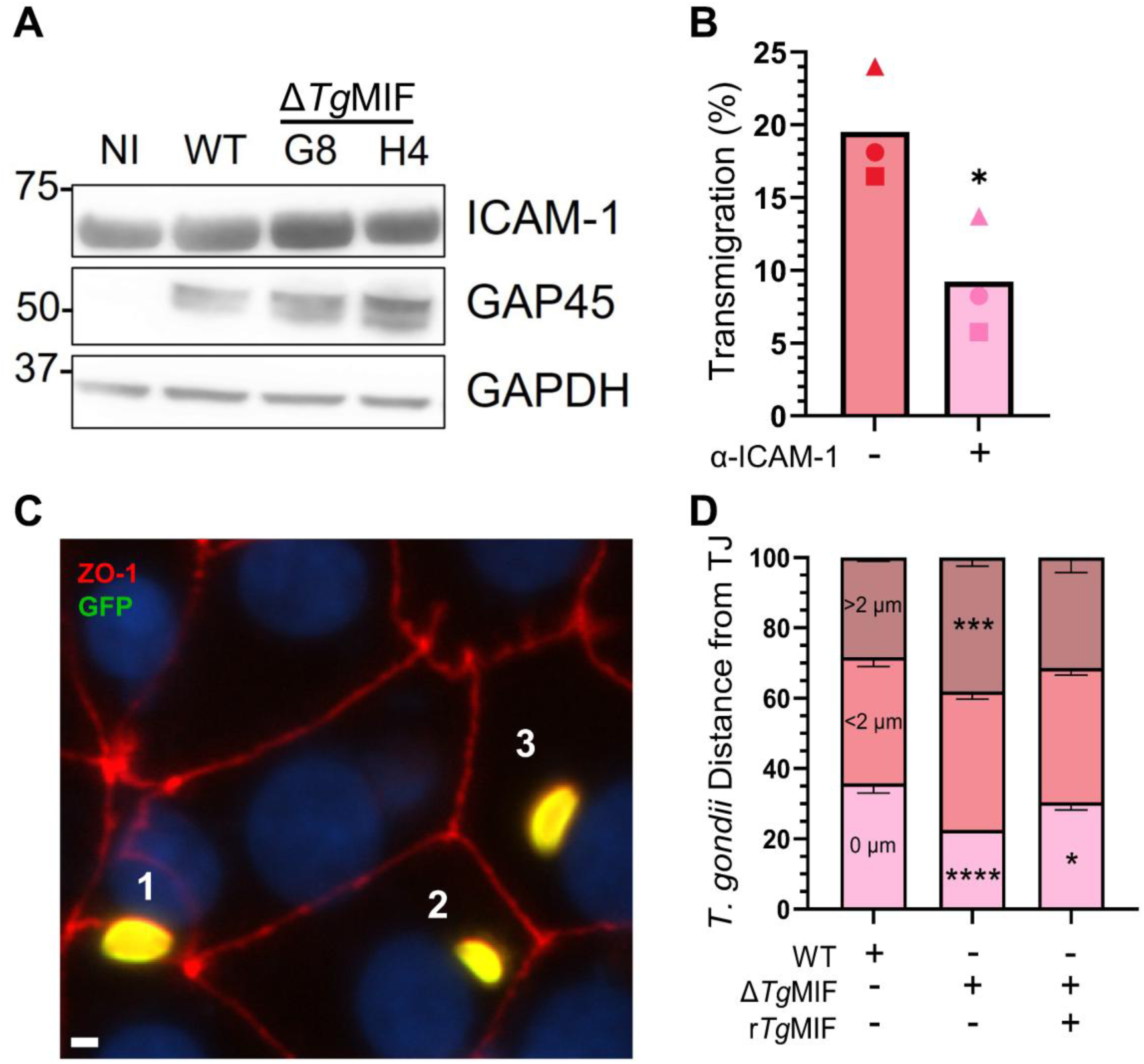
*Tg*MIF mediates extracellular parasite localization at the cellular tight junctions. **(A)** SDS-PAGE analysis of lysates from placental barriers infected or not (NI) for 5 hours with WT (RH-Luc) or Δ*Tg*MIF parasite, blotted with anti-ICAM-1, anti-GAPDH (cell loading control), and anti-GAP45 (parasite loading control), as well as their respective secondary HRP-antibodies. **(B)** WT parasites transmigration across the placental barrier pre-treated for 6 hours or not with an ICAM-1 neutralizing antibody [5 μg/mL]. (*n=3*, means ± SD). Student’s t-test, \**P* = 0.0349. **(C)** Localization of extracellular WT and Δ*Tg*MIF parasites on the surface of the placental barrier, treated or not with r*Tg*MIF parasites, 5 hours after infection. The staining was performed with anti-ZO-1 (red), anti-GAP45 (green), and nucleus (blue). (1) represents parasites localized at the tight junction, (2) parasites within 2 μm from the tight junction, and (3) parasites at a distance greater than 2 μm from the tight junctions. (Scale bar, 2 μm). **(D)** Quantification of parasite localization as defined in (C), for WT and Δ*Tg*MIF parasites. (*n=3*, 300 random parasites each, means ± SD). Two-way ANOVA with Dunnett’s multiple comparisons, ns (not significant), \**P* = 0.0306, \*\*\**P* = 0.0002, \*\*\*\**P* < 0.0001.

### *Tg*MIF is a cytosolic protein, excreted by the extracellular parasite via the ABC transporter

Even though r*Tg*MIF is functionally active in this study and previous ones ^37^, *Tg*MIF lacks a signal peptide, which is required for its secretion via typical secretory organelles ^37,57^. However, it is possible that, as *h*MIF ^58^ or *Entamoeba histolytica* MIF ^59^, *Tg*MIF is also excreted via a non-classical ATP-binding cassette (ABC) transporter and could be present in the parasite excretory/secretory antigens (ESA). Classical microneme and dense granule proteins are also found in the ESA ^60,61^. To test this, we first determine the subcellular localization of *Tg*MIF within the parasite, in a strain where the protein is endogenously HA-tagged using the pLIC plasmid ^62^. As expected, like *h*MIF, *Tg*MIF predominantly localizes within the cytosolic compartment of the parasite (**Figure 5A**). Then, we incubated *Tg*MIF-HA-tagged extracellular parasites in PBS/FBS (no detergent) at 37 °C for 3 hours, with or without probenecid, brefeldin A, or DMSO. Probenecid is an ABC transporter inhibitor, while brefeldin A disrupts the ER-Golgi transport ^59,61^. ESA was obtained after centrifugation at 18,000 xg for 30 minutes at 4 °C. Importantly, the PBS/FBS incubation did not compromise the integrity of the parasite plasma membrane, as demonstrated by propidium iodide staining (**Figure S5A**). Similar to GRA5 and MIC2 ^60,61^, *Tg*MIF is also present in the ESA fraction (**Figure 5B**). To investigate the impact of probenecid on *Tg*MIF excretion, we quantified the residual protein remaining in the parasite pellet and utilized GAP45 as a loading control. Our findings revealed a non-significant increase in *Tg*MIF retention within the parasite pellet during probenecid and probenecid/BFA treatments (**Figure 5C**), indicating that probenecid reduces the excretion of this protein into the ESA. Next, we hypothesize that decreased *Tg*MIF excretion will reduce WT parasite transmigration. We treated the WT parasite with probenecid or DMSO, or left it untreated, during transmigration. Probenecid, DMSO, or no treatment was added to the plaque assay to determine the parasite viability, which was similar between all conditions. We found that in the presence of probenecid, parasite transmigration decreased significantly compared to DMSO- or non-treated parasites (**Figure 5D**). Furthermore, to confirm that *Tg*MIF functions as an excreted effector, we conducted experiments in which increased concentrations of ESA from WT (WT-ESA) or Δ*Tg*MIF (Δ*Tg*MIF-ESA) were added to transmigrating WT or Δ*Tg*MIF parasites. In a concentration-dependent manner, the addition of WT-ESA significantly enhanced the transmigration of both WT and Δ*Tg*MIF parasites (**Figures 5E and F**). In contrast, Δ*Tg*MIF-ESA did not induce a similar effect (**Figures 5E and F**). Finally, r*Tg*MIF but not LPS, significantly enhanced transmigration in both WT and Δ*Tg*MIF parasites (**Figure 5G**). Thus, we clearly demonstrated that extracellular parasites excrete the cytosolic *Tg*MIF protein as a soluble factor to mediate transmigration.

**Figure 5.**
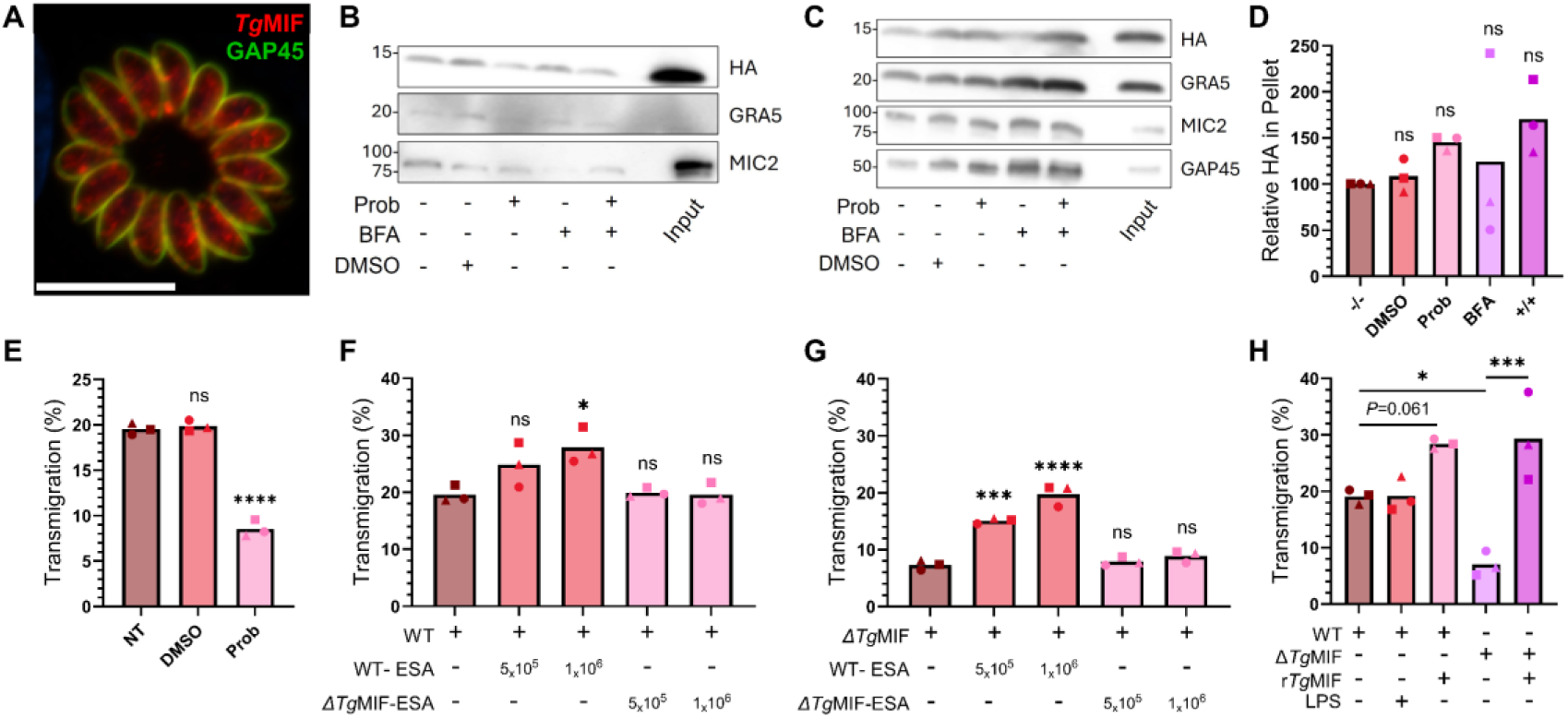
*Tg*MIF is a soluble, excreted effector. **(A)** Intracellular localization of *Tg*MIF-HA within tachyzoites after 48 hours of infection in HFF. Staining with anti-HA (red) and anti-GAP45 (green) antibodies. (Scale bar, 10 μm). **(B-C)** SDS-PAGE analysis of extracellular *Tg*MIF-HA-tagged parasites incubated for 3 hours in PBS/FBS at 37 °C in the presence or absence of Probenecid (Prob) [10 μM], Brefeldin A (BFA) [10 μM], or DMSO vehicle. For (B), ESA was TCA precipitated, and in (C), the parasite pellet was lysed. Input = 10% non-treated parasite pellet. The blotting was performed with anti-HA, anti-GRA5, anti-MIC2, and anti-GAP45 antibodies, with anti-GAP45 as a parasite loading control. **(D)** Relative quantification of *Tg*MIF-HA left in the parasite pellet of each condition from (C) after normalization with GAP45. (*n=3*, means ± SD). One-way ANOVA with Dunnett’s multiple comparisons, ns (not significant). **(E)** Transmigration across the placental barrier of WT parasites treated or not (NT) with Prob [10 μM] or DMSO vehicle. (*n=3*, means ± SD). One-way ANOVA with Dunnett’s multiple comparisons, ns (not significant), \*\*\*\**P* < 0.0001. **(F)** WT parasites transmigration across the placental barrier with or without ESA from WT or Δ*Tg*MIF parasites (equivalent of 5×10^5^ and 1×10^6^ parasites). (*n=3*, means ± SD). One-way ANOVA with Dunnett’s multiple comparisons, ns (not significant), \**P* = 0.0226. **(G)** Δ*Tg*MIF parasites transmigration across the placental barrier with or without ESA from WT or Δ*Tg*MIF parasites (equivalent of 5×10^5^ and 1×10^6^ parasites). (*n=3*, means ± SD). One-way ANOVA with Dunnett’s multiple comparisons, ns (not significant), \*\*\**P* = 0.0006, \*\*\*\**P* < 0.0001. **(H)** WT and Δ*Tg*MIF parasites transmigration across the placental barrier in the absence or presence of 100 ng/mL r*Tg*MIF or 100 ng/mL LPS. (*n=3*, means ± SD). One-way ANOVA with Sidak’s multiple comparisons, ns (not significant), \**P* = 0.0149, \*\*\**P* = 0.0002.

### *Tg*MIF is a factor determining *T. gondii* strain differences in transmigration capacity

Previous studies have demonstrated variations in the transmigration capacity among different strains ^28^. Based on this observation, we hypothesize that *Tg*MIF mediates these strain differences through its expression level, as no protein sequence differences were observed across strains (ToxoDB data). To test our hypothesis, we quantified *Tg*MIF expression levels by qPCR across seven archetypal RH-Luc and its parental strains RH88, ME49, and non-archetypal strains from South America, FOU, COUGAR, ARI, and GUY-DOS ^25,26^. The Δ*Tg*MIF strain was used as a reference, as it does not express *Tg*MIF. At the same time, we determined the transmigration capacity of each strain (normalized to their viability) using our placental barrier. We observed a positive correlation between *Tg*MIF expression levels and the transmigration capacity of strains, thereby supporting our hypothesis (**Figure 6A**). The alignment of the promoter regions among these strains shows that FOU, which exhibited the lowest *Tg*MIF expression and transmigration capacity, differs from the other strains. In contrast, COUGAR and ARI, who demonstrated the highest transmigration capacities, possess closely related promoter sequences (**Figure 6B**). Thus, we confirmed that the expression level of *Tg*MIF is a strain-determining factor that mediates the capacity of *T. gondii* strains to transmigrate across the human *in vitro* placental barrier.

**Figure 6.**
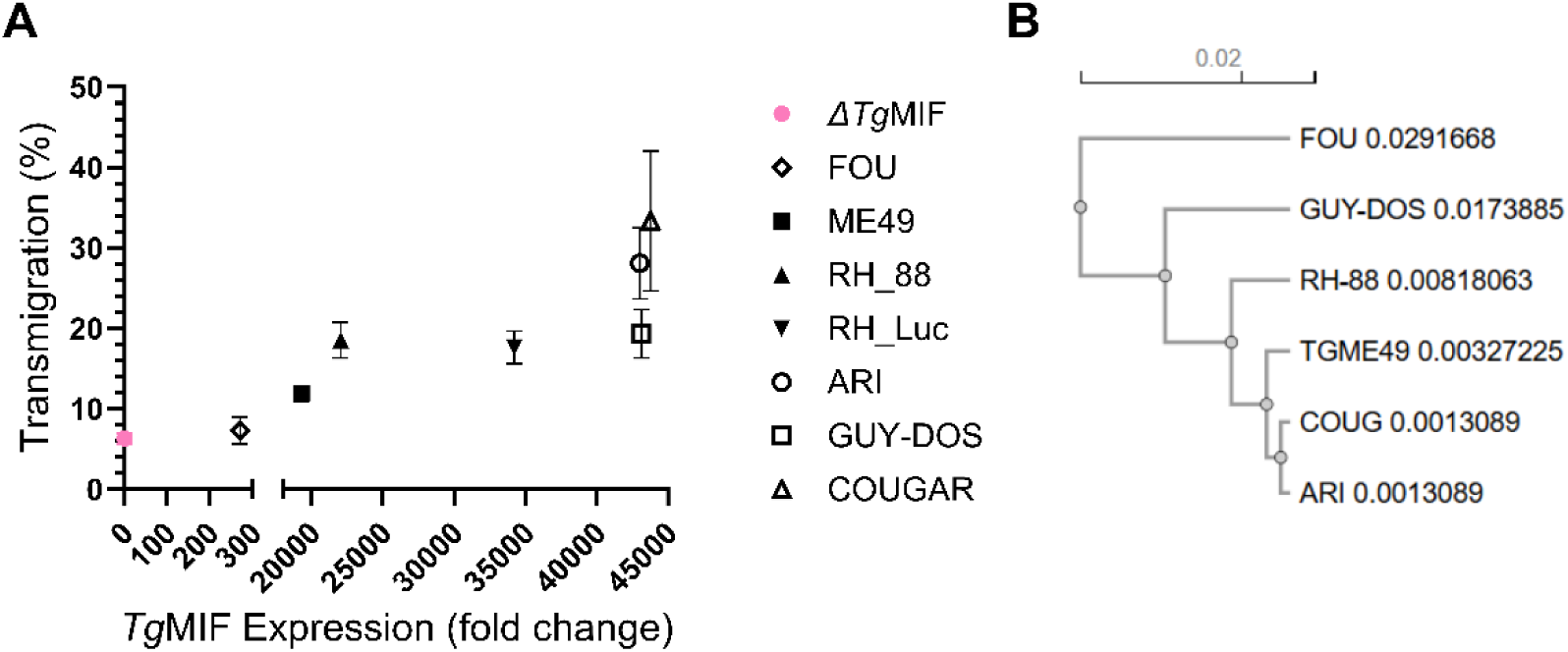
*Tg*MIF expression level determines the transmigration capacity of *T. gondii strains*. **(A)** Positive correlation between *T. gondii* strain transmigration capacity (y-axis) and *Tg*MIF mRNA expression level in each strain (x-axis). *Tg*MIF mRNA level in each strain was normalized to TubA1 level, and Δ*Tg*MIF serves as the reference strain. (*n=3*, means ± SD). Spearman correlation, r = 0.9524, *P* = 0.001. **(B)** Guide tree depicting the relatedness of the strains based on the alignment of their promoter sequence using Clustal Omega.

## DISCUSSION

Very few studies have investigated how *T. gondii* crosses the placental barrier, possibly because it is perceived as a random occurrence or due to the scarcity of accurate placental models. Using *h*TSCs ^20,39^, we have developed a human *in vitro* placental barrier that demonstrates significant resistance to the intracellular replication of *T. gondii*. This resistance is likely attributed to cell polarization and the presence of extensive areas of STBs, which were absent in the previous model using the BeWo cell line ^31^. The *h*TSCs have been used to create a barrier consisting of a layer of STBs situated above progenitor CTBs ^40^. Although this barrier represents a more robust structural model compared to ours, quantifying transmigration would pose a challenge. This is due to the limited number of intercellular junctions, which would necessitate infecting the barrier with a substantially larger number of parasites. In fact, the exact number of parasites capable of successfully traversing the placenta to infect the fetus during pregnancy remains unknown. This is likely a low number, as the number of parasites that reach the placenta would be constrained by factors limiting dissemination. Using our model, we demonstrate that across various parasite strains, the number of extracellular parasites that transmigrate is relatively low, resulting in minimal disruption of placental barrier integrity. Therefore, extracellular parasites’ transmigration could explain why up to 70% of fetal infections are discreet phenomena that go undetected during pregnancy ^63,64^. Through our system, we have demonstrated that extracellular parasite transmigration is not a matter of chance, but rather an active process mediated by a parasitic effector.

Our findings reveal that *Tg*MIF is localized within the parasite cytosol and shows no colocalization with the classical secretory organelles. Lacking a signal peptide, *Tg*MIF, similar to host and several parasitic MIFs, is excreted through the ABC transporter pathway ^58,59^ into the ESA fraction ^59^. While *h*MIF is excreted by ABCA1 ^58^, which is absent in *T. gondii*, the parasite still expresses a diverse array of ABC transporter families, many of which remain uncharacterized ^65,66^. Classical secreted microneme and dense granule proteins are also found in the ESA ^60,61,67^. However, the excretion of GRAs into the ESA is influenced by serum, temperature, and pH ^61^, whereas the release of MICs is mediated by calcium signaling ^67^. Notably, ESA has been extensively utilized in various *T. gondii* studies, including immunization strategies for vaccine development ^68^, modulation of the host immune response ^69,70^, and the mediation of miscarriage in mice via Toll-like receptor 4 ^71^. However, the molecular factors mediating these ESA functions have yet to be identified. In this study, we show that *Tg*MIF, present in the extracellular parasite ESA, plays a crucial role in modulating the placental barrier during *T. gondii* transmigration. Our findings emphasize the role of excretory proteins in modulating host pathways before infection, which could also affect how classical secreted proteins subsequently interact with the host.

*In vitro*, r*Tg*MIF binds to CD74 receptor ^38^; however, we found that *Tg*MIF does not mediate transmigration via CD74 and does not require *h*MIF. To transduce the signal from the ligand, CD74 requires distinct coreceptors, including CD44, CXCR2, CXCR4, and CXCR7 ^72,73^. However, some of these coreceptors can interact with the ligand independently of CD74 ^73,74^. For example, *Pf*MIF interacts with CXCR2 and CXCR4 independently of CD74 to inhibit the random migration of monocytes ^75^. Similarly, our characterization of *Tg*MIF function supports the possibility of binding to two distinct receptors. We show that *Tg*MIF activates ERK1/2 MAPK within 45 minutes and induces FAK dephosphorylation within 5 hours. Both ERK1/2 MAPK activation and FAK dephosphorylation are critical for parasite transmigration. However, our data exclude the possibility that FAK dephosphorylation is dependent on the ERK1/2 MAPK pathway via PIN/PEST phosphatases activation ^51^. Therefore, FAK dephosphorylation will affect ZO-1 at the tight junction ^32,76^, while ERK1/2 MAPK activation could direct the phosphatase PP2A to dephosphorylate and inactivate occludin ^77^, which we did not test in this study. Supporting literature on Caco-2 cells shows that infection with *T. gondii* results in reduced occludin expression and visible disruption observed by immunofluorescence ^78,79^. Thus, we established that *Tg*MIF mediates extracellular parasite transmigration in a CD74-independent manner, highlighting its functional specificity compared to *h*MIF.

Our research shows that *Tg*MIF mediates the localization of extracellular parasites at cellular tight junctions and confirms that the host adhesion molecule ICAM-1 is essential for their transmigration ^31^. Interestingly, *T. gondii* does not induce the upregulation of ICAM-1 expression at the placental barrier, as this was observed with *h*MIF ^52^. Thus, both *Tg*MIF and ICAM-1 could mediate transmigration by enhancing the adhesion of extracellular parasites to cellular tight junctions. During leukocyte transmigration, ICAM-1 binding to its ligand can activate the ERK1/2 MAPK pathway, thereby stimulating more ICAM-1 expression ^45^. However, it is noteworthy that r*Tg*MIF alone can activate ERK1/2 MAPK, suggesting that prior contact between the parasite and the cell membrane via ICAM-1 is not necessary for this activation. In one scenario, the activation of ERK1/2 MAPK by *Tg*MIF results in the upregulation of other adhesion molecules, such as VCAM-1, which, together with ICAM-1, facilitates transmigration ^45,80–82^. Conversely, in another scenario, *Tg*MIF will only induce the relocalization of ICAM-1 to cellular tight junctions without altering its expression level ^83^. Both scenarios are plausible, as in our data, the inhibition of ICAM-1 or the absence of *Tg*MIF produces similar effects on parasite transmigration. Thus, our study suggests that *Tg*MIF mediates the adhesion of extracellular parasites to tight junctions before facilitating transmigration. Therefore, it will be interesting to study how parasite motility changes upon contact with the cellular tight junction.

We found that *Tg*MIF expression levels mediate strain differences in transmigration capacity. A previous study also showed differences in transmigration capacity between archetypal strains ^28^. In this study, the RH type I strain presents the highest transmigration capacity, which is supposed to be mediated by the LDM ^28^. The molecular factors behind the LDM phenotype are unknown; however, it not only mediates transmigration but also virulence, migration, and host cell metabolic exploitation ^28,84^. It is possible that *Tg*MIF, through a synergistic effect with other effectors, mediates LDM; however, our study did not investigate this further. We found that RH transmigration capacity is higher than that of ME49, a type II strain; however, it is much lower than the non-archetypal strains COUGAR, GUY-DOS, and ARI. We clearly show that the transmigration capacity of a strain depends on *Tg*MIF expression levels in this strain. Indeed, the expression level of *Tg*MIF varies among *T. gondii* strains ^26^, and our data confirms this and provides a phenotypic outcome. However, a previous study showed that *Tg*MIF expression level is higher in ME49 than in RH ^85^. We extracted mRNA from intracellular parasites; we cannot rule out the possibility that, if the extraction had been performed on extracellular RH, transcript levels might differ from those in our study. The alignment of the *Tg*MIF promoter supports our data, where FOU has the most divergent sequence and the lowest expression and transmigration. At the same time, COUGAR and ARI are similar and display high expression and transmigration. Finally, we did not test this, but it is possible that the excretion levels of *Tg*MIF in COUGAR, ARI, and GUY-DOS account for the differences in their transmigration capacity. Thus, our data suggest that parasite strains with high *Tg*MIF expression and excretion are most likely to cause congenital infection in pregnant women.

In conclusion, we propose the model shown in Figure 7, in which extracellular parasites release *Tg*MIF, which interacts with two distinct receptors on the maternal side of placental barrier cells. Binding to Receptor 1 activates the ERK1/2 MAPK pathway, resulting in the modulation of cell adhesion molecules (CAMs) at the tight junctions and/or the dephosphorylation of Occludin. As a result, the adherence of extracellular parasites to the placental barrier increases, particularly at the cellular tight junctions. Simultaneously, binding to Receptor 2 leads to FAK dephosphorylation and affects the localization of ZO-1 at the tight junctions. These modulation enables the parasites to transmigrate across the permeable junction and reach the fetal side.

**Figure 7.**
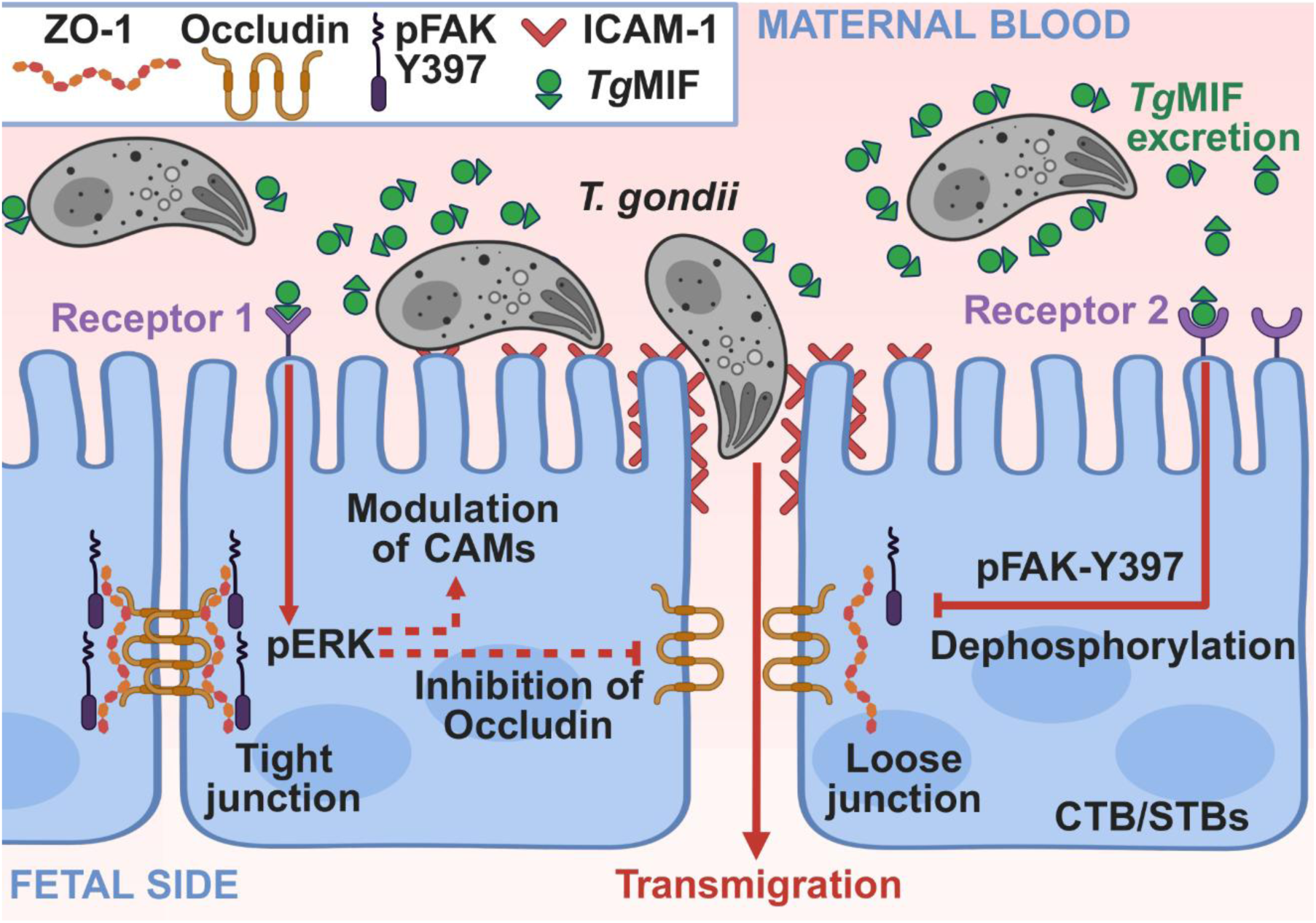
Model of *Tg*MIF mediating *T. gondii* transmigration across the placental barrier. Extracellular parasites on the maternal side of the barrier excrete *Tg*MIF, which then interacts with unknown receptors 1 and 2. Via receptor 1 and the ERK MAP1 pathway, *Tg*MIF mediates modulation of ICAMs and/or occludin proteins. Via receptor 2, *Tg*MIF mediates the dephosphorylation of FAK. Together, increased parasite adhesion to tight junctions and loosening of junctions facilitate transmigration toward the fetal side of the barrier.

## MATERIALS AND METHODS

### Parasite Culture

*T. gondii* parasites included in this study are the RH-Luc^+^ (WT) ^41^, RH-Luc^+^_Δ*Tg*MIF (Δ*Tg*MIF), RH-Luc^+^_Δ*Tg*MIF_*h*MIF-HA, (*h*MIF comp) RH-Luc^+^_Δ*Tg*MIF_*Tg*MIF-HA (*Tg*MIF comp), RHΔku80_*Tg*MIF-HA, RH-88, ME49, ARI, COUGAR, FOU, and GUY-DOS ^26^. All parasites were passaged on a monolayer of human foreskin fibroblasts (HFFs) growing in Dulbecco’s Modified Eagle Medium (Gibco, #11965118) supplemented with 1% Heat heat-inactivated fetal Bovine Serum (Gibco, #A56698-01), 100 U/mL Penicillin-Streptomycin (Gibco, #15140122), 2 mM L-Glutamine (Gibco, #25030081), and 10 μg/mL Gentamicin (Gibco, #15710072) at 37°C in 5% CO_2_.

### Generation of an *in vitro* human placental barrier

To prepare the *in vitro* human placental barrier, 12-well transwells with 8 μm pores (Falcon #353182) were pretreated for 2 hours with 50 μg/mL human placental collagen IV (MilliporeSigma #C5533-5MG), then for 15 minutes with iMatrix-511. *h*STC in CTB media ^20^ were seeded at a cell density of 3 ×10^4^ cells/transwell and cultured for 8-10 days until reaching confluency, with media changes every 48 hours. As a control for non-barrier-forming cells, the above was repeated using HFFs and their respective media. Transepithelial electrical resistance was measured using an ohmmeter (World Precision Instruments, #EVOM3) blanked with a sterile transwell in fresh media. A threshold of 150 Ω*cm² was established for the polarized placental barrier in future experiments. Barrier integrity was measured by adding 20 μg/mL of 40kDa FITC-Dextran (MilliporeSigma #FD40S-250MG) to the apical side of the transwell. Sixteen hours later, the media on the basolateral side of the transwell was collected. FITC-Dextran passage was quantified using a SpectraMax iD3 (Molecular Devices) with an excitation λ of 485 nm and an emission λ of 528 nm. A sterile transwell with no cell culture was included as a blank to determine the maximum of passing FITC-Dextran. FITC-Dextran passage is then presented as the percentage of FITC-Dextran measured in a sample compared to the blank. Media control was included during measurements and subtracted before other calculations. For all transmigration, parasite growth, and luciferase assays, the A83-01 inhibitor was removed, and the barrier was washed 24 hours prior ^20^.

### Plasmid Generation

sgRNAs targeting the *Tg*MIF gene (TGGT1_290040) were cloned into the pU6-Universal vector ^86^. For C-terminal HA epitope tagging, a region of the *Tg*MIF gene upstream of the stop codon was amplified by PCR with specific primers and inserted into pLIC-HA-dhfr using ligation-independent cloning ^62^. For the recombinant *Tg*MIF plasmid, the *Tg*MIF coding sequence was amplified with specific primers from wild-type RH parasites’ cDNA and then inserted into the pET21a+ plasmid (Addgene #69740-3) at the NdeI and XhoI cut sites. For complementation plasmids, *Tg*MIF and *h*MIF coding sequences were amplified with specific primers (the reverse primer included an HA epitope) from wild-type RH parasites and HFF, respectively. Both were then inserted into pUPRT::DHFR-D plasmid (Addgene #58528) ^87^ using Gibson Assemble, flanked by 1000 base pairs upstream (promoter) and downstream (3’UTR) of the *Tg*MIF gene. The promoter was amplified from DNA synthesized by Integrated DNA Technologies; the full-length sequence is available in Supporting Information. All sgRNAs and primer sequences are listed in the Primer Table of Supporting Information.

### Construction of Parasite Strains

To generate the Δ*Tg*MIF strain, the plasmid containing sgRNAs was co-transfected with NotI-linearized pTKOatt, which includes the *hxgprt* selection cassette and GFP ^88^, into RH-Luc^+^ (parasites at a 5:1 ratio of sgRNA to linearized pTKOatt plasmid). 24 h post-transfection, two distinct populations were selected with mycophenolic acid (50 μg/mL) and xanthine (50 μg/mL) and cloned by limiting dilution. PCR and sequencing were used to confirm two distinct individual knockout clones, Δ*Tg*MIF (G8 and H4). Complementation with *Tg*MIF and *h*MIF genes was performed in the Δ*Tg*MIF strain by transfecting with either pUPRT-*Tg*MIF-HA or pUPRT-*h*MIF-HA plasmid. After first lyse out, populations were selected with 10 μM 5-fluoro-2’-deoxyuridine (FUDR) (Millipore Sigma #F0503-100MG) and cloned by limiting dilution. Complemented parasites were isolated after HA immunofluorescence assay. Endogenously tagged parasites were made in the RHΔku80 strain ^62^ by transfection with plasmid pLIC-*Tg*MIF-HA-dhfr. 24 h post-transfection, populations were selected with 1 μM of pyrimethamine and cloned by limiting dilution. Clones were isolated after an HA immunofluorescence assay.

### Recombinant Protein Production

The plasmid pET21a+ containing *Tg*MIF gene coding sequence was used to transform BL21(DE3) competent E. coli (New England BioLabs #C2527H). *Tg*MIF-6xHis protein expression was induced using 1 mM IPTG at 37°C for 16 hours. The E. coli was centrifuged down and sonicated on ice for 30 seconds on, 60 seconds off, and 20 times in PBS containing lysozyme and DNaseI. Recombinant *Tg*MIF-6xHis protein (r*Tg*MIF) was recovered using NEBExpress Ni-NTA Magnetic Beads (New England BioLabs #S1423S) according to the manufacturer’s protocol, eluted with 0.5 M imidazole, dialyzed and cleaned of endotoxins using the Pierce High Capacity Endotoxin Removal Spin Columns (Thermo Scientific #88274) according to the manufacturer’s protocol.

### Immunofluorescence Assay

All placental barriers or HFFs, infected or not, were fixed for 20 minutes at room temperature with 4% Paraformaldehyde, then blocked and permeabilized for 30 minutes with 3% Bovine Serum Albumin in PBS with 0.2% Triton X-100. Respective primary antibodies were used, as well as the following Alexa-Fluor secondary antibodies [1:1,000] (Invitrogen, α-Mouse-594 #A11005, α-Rat-594 #A11007, α-Rabbit-488 #A11008, α-Rabbit-594 #A11012). HOECHST [1:2,000] (MilliporeSigma #63493-5MG) was used to visualize the nucleus. All images were taken at 40x or 100x objective on a Nikon ECLIPSE Ti2-A. **For placental barrier validation**, the cells were stained with Rat α-ZO-1 [1:1,000] (Cell Signaling Technology #6B6E4) overnight at 4°C (rocking), and Rabbit α-Syndecan-1 [2 μg/mL] (Abcam #AB128936) at room temperature for 1 hour. **For parasite per vacuole assay,** WT and Δ*Tg*MIF-infected placental barriers or HFFs at the multiplicity of infection 1 (MOI1). The cells were then stained with Rabbit α-GAP45 [1:5,000] antibody. Five randomized images were taken at 40x, utilizing a 10-step Z-Stack for the placental barrier due to cell thickness. From these images, all parasite vacuoles were quantified for the number of parasites within the vacuole and grouped into 1, 2, 4, 8, or 8+ parasites per vacuole. **For the tight junction proximity assay**, Placental barriers were treated with 100 ng/mL r*Tg*MIF or left untreated, then infected with 2×10^5^ WT or Δ*Tg*MIF parasites. The barriers were incubated for 5 hours and stained with ZO-1 and GAP45 antibodies. Images of each well were taken at a 40x objective, and a total number of 300 parasites was considered. The NIS-Elements tool was used to measure parasites’ proximity to the cellular tight junction, with distances of 2 μm or less considered closer. **For *Tg*MIF intracellular localization**, HFFs infected with RHΔku80_*Tg*MIF-HA for 24 hours at MOI1 were stained with Rat α-HA antibody (Sigma Aldrich, #NC1821908) for 1 hour at room temperature, and a representative image was captured at 100x to display *Tg*MIF intracellular localization.

### Luciferase Assay

The placental barrier and HFF were infected with RH*Δhpt*_Luc^+^ (WT) for 24 hours. Cells were collected and lysed at 37°C for 10 minutes in 75 μL Cell Lysis Buffer ^89^ made of PBS containing 10% glycerol, 1% Triton X-100 (MilliporeSigma #TX1568-1), 0.2% dithiothreitol (MP Biomedicals #100597), and protease inhibitor (Thermo Scientific #1861284). 25 μL from each sample was plated in technical triplicate in a 96-well plate and then measured on the SpectraMax iD3 after automatic injection of click-beetle luciferin (Promega #E1603). Data are shown as the mean of each biological replicate, with uninfected cell control values subtracted.

### Excreted Secreted Antigen Collection

T175 flasks were infected with 1×10^7^ of freshly lysed WT, Δ*Tg*MIF, or RHΔku80_*Tg*MIF-HA parasite for 3 days. Freshly lysed parasites were collected from their respective flasks and centrifuged at 582 × g for 10 minutes. The supernatant was aspirated, and the parasites were resuspended in 5 mL PBS. Parasites were counted, and fractions of 5×10^7^ were centrifuged at 582 x g for 10 minutes. Each fraction was resuspended in 1 mL PBS + 10% FBS and then rested at 37°C for 3-5 hours in the presence or absence of 10 μM Probenecid (MedChemExpress #HY-B0545), 10 μM Brefeldin A (MedChemExpress #HY-16592), or an equivalent volume of DMSO. Excreted Secreted Antigen (ESA) samples from WT and Δ*Tg*MIF parasite strains used for transmigration assays were centrifuged at 18000 x g, 4°C for 30 minutes to separate parasites from supernatant, with supernatant then stored at −80°C until use. The ESA from the RHΔku80_*Tg*MIF-HA parasite was isolated similarly, precipitated with Trichloroacetic Acid (Millipore Sigma #T6399), and resuspended in PBS. The RHΔku80_*Tg*MIF-HA parasite pellets in RIPA buffer and the corresponding resuspended ESA were both processed for Western blot analysis as described below.

### SDS-PAGE and Western Blot

Placental barriers were treated with 50 μM PD98059 or an equivalent volume of DMSO for 24 hours, then infected or not for 45 minutes. All Western blot samples were processed as follows. Cells were collected and lysed in RIPA buffer on ice for 30 minutes, then centrifuged at 18000 xg, 4°C for 30 minutes. The supernatant was collected and combined 1:4 with 4x loading buffer containing 375 mM Tris-HCl, 50% glycerol, 10% SDS, and 0.03% Bromophenol Blue. Samples were then boiled at 50°C for 5 minutes (for the detection of phosphorylated protein) or at 90°C for other samples. Samples were run on 10-12% SDS-PAGE and then transferred to a PVDF membrane. The membrane was blocked in 5% milk in Tris-buffered saline containing 1% Tween 20. GAPDH [1:1,000] (Cell Signaling Technology #D16H11) and GAP45 [1:3,000] antibodies were used for cell and parasite loading control, respectively. All quantification was normalized to GAPDH. Primary antibodies were added at the following concentrations: Rat α-HA [1:500], Mouse α-ICAM-1 [1:250] (Invitrogen, #ENMA5407), Rabbit α-pERK1/2-T202/Y204 [1:2,000] (Cell Signaling Technology #D13.14.4e), Rabbit α-total ERK1/2 [1:1,000] (Cell Signaling Technology #137F5), Rabbit α-pFAK-Y397 [1:750] (Cell Signaling Technology #3283), Rabbit α-total FAK [1:1,000] (Cell Signaling Technology #3285T), Mouse α-GRA5 [1:5,000] (BioVision #A1299-50), and Rabbit α-MIC2 [1:5,000]. After incubation with respective secondary HRP antibodies, protein visualization was performed by chemiluminescence using ProSignal Dura (Genesee Scientific #20-301B).

### Transmigration Assay

Freshly lysed parasites were seeded onto the placental barrier at 2×10^5^ parasites/transwell, then left untouched at 37°C, 5% CO_2_ for 16 hours. From the same parasite population, three wells of confluent HFF cells in a 24-well plate were infected with 100 parasites and left untouched at 37°C and 5% CO_2_ for 5 days. One placental barrier was left uninfected as a control. To compare across several strains in **Figure 6**, the initial infection was increased to 5×10^5^ parasites/transwell to account for low viability in some strains. After 16 hours, media from the basolateral side of the transwells was collected and centrifuged at 582 xg for 10 minutes. The supernatant was removed, leaving 100 μL to resuspend the transmigrating parasites, which were then counted on a hemocytometer. After 5 days, the number of plaques in each well of the 24-well plate was counted. The average number of plaques was considered equivalent to parasite viability. The number of transmigrating parasites was normalized to this viability, yielding the percentage of transmigrating viable parasites. Treatment of placental barrier with inhibitors and proteins included the following: 500 ng/mL recombinant HIV-Nef protein (Abcam, #ab63996) concurrent with infection, 100 ng/mL of human IFN-γ (Peprotech #300-02-200UG) added 6 hours before infection, 100 ng/mL *E. coli* K12 lipopolysaccharide (InvivoGen #tlr-eklps) concurrent with infection, 100 ng/mL r*Tg*MIF protein concurrent with infection, 5 μg/mL of ICAM-1 neutralizing antibody (Invitrogen #ENMA5407) 6 hours before infection, 50 μM MEK inhibitor PD98059 (Cell Signaling Technology, #9900S) or an equivalent volume of DMSO 24 hours before infection, 10 μM human MIF inhibitor ISO-1 (Sigma-Aldrich #475837-5MG) 30 minutes before infection, 12.5 μM FAK inhibitor PF-573228 (Sigma-Alrich #PZ0117-5MG) 16 hours before infection, 10-50 μg/mL Milatuzumab (MedChemExpress #HY-P99731) 24 hours before infection, 10 μM Probenecid or an equivalent volume of DMSO for 10 minutes before infection, a volume of WT or *ΔTg*MIF Excreted-Secreted Antigen (ESA) equal to 5×10^5^ or 1×10^6^ parasites concurrent with infection.

### Mouse Survival Assay

Mice were housed in an Association for Assessment and Accreditation of Laboratory Animal Care International-approved facility at Texas A&M University. All animal studies were conducted in accordance with the US Public Health Service Policy on Humane Care and Use of Laboratory Animals, and the Institutional Animal Care and Use Committee at Texas A&M University in College Station approved protocols. Eighteen 8-week-old CD-1 mice (Charles River Laboratories) were infected intraperitoneally with a lethal dose of 2×10^3^ WT or Δ*Tg*MIF parasites, 9 mice per group. Mice were observed twice daily for signs of infection and weighed once every two days. Survival was recorded daily until all mice had died or were euthanized.

### qPCR

mRNA was isolated from intracellular parasites using Monarch® Total RNA Miniprep Kit (NEB #T2110), and cDNA was obtained after RT-PCR using LunaScript® RT SuperMix (NEB # M3010L). For each parasite strain, *Tg*MIF transcript levels were quantified by qPCR using specific primers (see the Primer Table in the Supporting Information). The transcript level of the *T. gondii* TUB1A gene was used to normalize the number of parasites across the strain, and RH/hpt^+^_Luc^+^_Δ*tgmif*_GFP^+^ (Δ*Tg*MIF) was used as a reference strain.

### Statistical Analysis

Statistical analysis for all experiments was performed in GraphPad PRISM, with the number of replicates, the statistical test used, and the significance values reported in the respective figure legend.

## ACKNOWLEDGMENTS

We thank Dr. Jon Boyle (University of Pittsburgh) for providing the *h*TSC and culture protocols. We thank Dr. Dominic Soldati (University of Geneva) for providing GAP45 antibody and Dr. Vern Carruthers (University of Michigan) for providing MIC2 and MIC5 antibodies. We thank Dr. Jeroen Saeij (University of California, Davis) for providing non-archetypal parasite strains. This work was supported by NIH R21 AI185518-02 grant.

## AUTHOR CONTRIBUTIONS

K.B.K. and L.O.S. designed research; K.B.K. and G.D.S. prepared parasite strains and recombinant protein; K.B.K. and L.O.S. performed experiments; K.B.K. and L.O.S. analyzed data; K.B.K. designed figures; and K.B.K. and L.O.S. wrote the manuscript.

