## Supplemental Figure and Table for "*Toxoplasma gondii Tg*MIF mediates the transmigration of extracellular parasites across the human placental barrier"

**SUPPLEMENTAL INFORMATION**

Synthesized *Tg*MIF Promoter Sequence
5’-
GACCTGTCCACAGGGCTTCTCACAAGTCCTTCGCCTTCTGCTTCCTTTTGATCTCGCGTTTTCTACAG
TTTTTCGCGCCCCTGTCTGGCGAGAGTAACCTCACATATAGGGCGTCTGGTTTTGTGCCTCTCCGAA
ACGGCCTCTCTGTCTCCGTACCTTTGCGCTCCTCCGCTGATAGAACGCATGCGCTGTTGCCGCTCG
GAGCTTTTTCCGTCGGTGGCTGGTTTGGTTGTTGAGACACAACTCCTCGCAACTCTCTACAGAAGCG
GAAGAAAAAGCCGAAGAAAAGACGAAGAATTGAGATCCTGAAGCGGTCTGGATCTCTGTCTTTCTCT
GTTTCCTCTTCACACGTCGCCACTTGTCCCCAATTCAACAAGAAGGCCTTCGGATTACCTGTCCGGA
GCGGTGTGCGAGTTACGCTGTCCACGGTCACTCCTGCGTCGCCGCAAATCTTCCCTATTTCTCGGTT
CTTTCTCTCTAAAGGTCTCTCCTCCGCTCGAAAAGCTCGTCTTCCTGGTGACCAGCAGAGAGCGACC
GTCTGCGTCGAACGGACGCCCGAGACAGCTTCTCTCGGAGCCGCGCCAGACGCAGAAGATTCTGG
CAGACTCGACCTCCCGCCTGTTGAGGCGCGCGTTTTCCGCGGGGGGGGGGGGTCGCATTTGCGGC
TCTTTCGATCGACACTAGAGTCTGGAGACACACAGACGCGAAGTCGAGCGTTTAGCAGCGCGTTTTT
GAGGCTGAAACAGAGGGAAAAGCGCGCTTGCGTCGAGACAAGCTGGCGCCGTTAACCTCACAGTC
GCCGCCTTCCCGTTTGAGCGAAGAAACAGCGAACGCGAGACGACTGAAATCCCGCGCTTTCTGCAC
AGACGGAGGCGCGCGCGCGAGTTGCGTCTCCGCCACCGGTGCACTCAGGAGTCCTCCGGAGAAAA
TTCACTTGGTGTCTTCACCCCTTTGCGTTCCGGGTCGCTCTGACTTTTTTTGTCTCTTTCTCTTCGTAC
ACCTTTCAAGTTCCCCCACAAA-3’

**Supplemental Table 1.** Primer Table.

| **Primer Name** | **Sequence** | **Description** |
|---|---|---|
| pET21_TgMIF_F | 5'-GGG GCA TAT GCC CAA GTG CAT GAT CTT TTG CC-3' | Forward primer for making r*Tg*MIF inserts for pET21a+ plasmid |
| pET21_TgMIF_R | 5'-GGG GCT CGA GGC CGA AAG TTC GGT CGC CCA TGG C-3' | Reverse primer for making r*Tg*MIF inserts for pET21a+ plasmid |
| sgRNA1_Tg290040_F | 5'-AAG TTG CAG CAG GAC GCC CTC TTG AG-3' | Forward primer for generating *Tg*MIF knockout sgRNA |
| sgRNA1_Tg290040_R | 5'-GAA GAA ACC GCG AAC GCG AGA CG-3' | Reverse primer for generating *Tg*MIF knockout sgRNA |
| pUPRT::GOI GIB F | 5'-GAC AGA CCG CTG ACG GAA TC-3' | Forward primer to amplify pURPT-DHFR without DHFR cassette |
| pUPRT::GOI GIB R | 5'-AGA AGC CCT GTG GAC AGG TC-3' | Reverse primer to amplify pURPT-DHFR without DHFR cassette |
| pUPRT prom290040 F3 | 5'-GAC CTG TCC ACA GGG CTT CTC-3' | Forward primer to amplify 1000 bp (upstream of ATG) of the promoter of the gene with pUPRT homology arm from synthesized DNA from IDT |
| pUPRT prom290040 R3 | 5'-TTT GTG GGG GAA CTT GAA AGG TG-3' | Reverse primer to amplify 1000 bp (upstream of ATG) of the promoter of the gene with pUPRT homology arm from synthesized DNA from IDT |
| pUPRT cDNA290040HA F | 5'-TAC ACC TTT CAA CCC CCA CAA AAT GCC CAA GTG CAT GAT CTT TTG CC-3' | Forward primer to amplify coding region of *tgmif* from cDNA |
| pUPRT CDNA290040HA R | 5'-TCA AGC GTA ATC TGG AAC ATC GTA TGG GTA GCC GAA AGT TCG GTC GCC CAT GGC CCA CTC-3' | Reverse primer to amplify coding region of *tgmif* from cDNA flanked with HA tag |
| pUPRT cDNA hMIF-HA F | 5'-TAC CAA TTT CAA GTT CCC CCA CAA AAT GCC GAT GTT CAT CGT AAA CAC CA-3' | Forward primer to amplify coding region of human MIF from cDNA |
| pUPRT cDNA hMIF-HA R | 5'-TCA AGC GTA ATC TGG AAC ATC GTA TGG GTA GGC GAA GGT GGA GTT GTT CCA GCC CAC ATT-3' | Reverse primer to amplify coding region of human MIF from cDNA flanked with HA tag |

| | | |
|---|---|---|
| pUPRT 3UTR290040 F | 5'-TAC CCA TAC GAT GTT CCA GAT TAC GCT TGA GCT GAG AGC GCT CTG GAC GTT TCA TGC AGA-3' | Forward primer to amplify 1000 bp of the 3' UTR of the gene |
| pUPRT 3UTR290040 R | 5'-GAT TCC GTC AGC GGT CTG TCT GTG AAA CTC TGT ATA TCT TTC TTG CTT TC-3' | Reverse primer to amplify 1000 bp of the 3' UTR of the gene with pUPRT homology arm |
| TUB1A Forward | 5'-GAC GAC GCC TTC AAC ACC TTC TTT-3' | Forward primer to amplify *Toxoplasma* TUB1A for qPCR as a parasite loading control |
| TUB1A Reverse | 5'-AGT TGT TCG CAG CAT CCT CTT TCC-3' | Reverse primer to amplify *Toxoplasma* TUB1A for qPCR as a parasite loading control |
| TgMIF Reverse_qPCR | 5'-ATG CGG TTC TTG GGG ACG CCC-3' | Reverse primer to amplify *tgmif* for qPCR of atypical strains |
| TgMIF Forward_qPCR Exo1 | 5'-CGC AGC AGG ACG CCC TCT TGA-3' | Forward primer to amplify *tgmif* for qPCR of atypical strains |

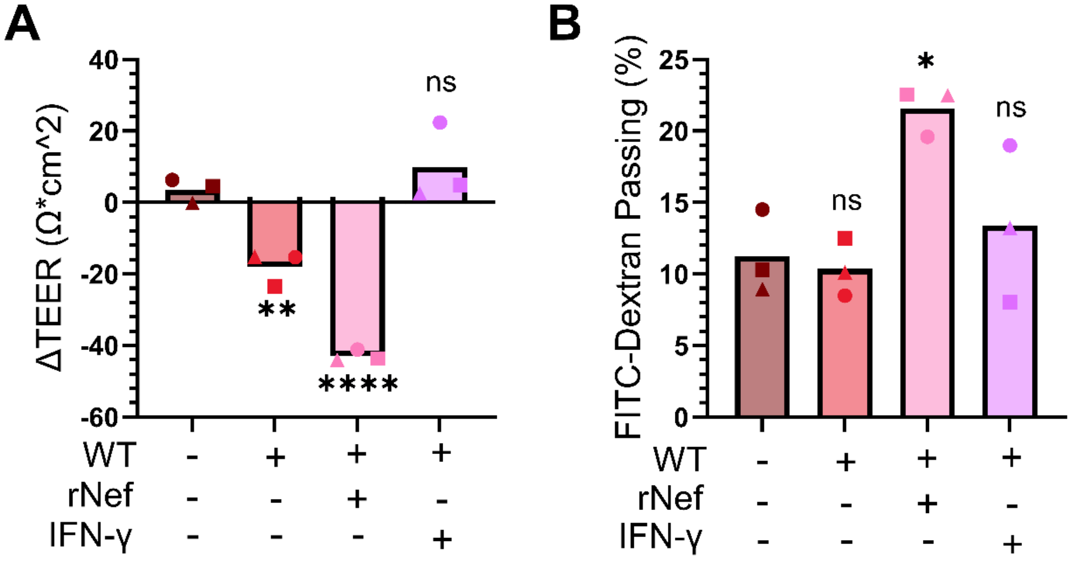


**Supplemental Figure 1. (A)** TEER measurement pre- vs post-transmigration, of RH-Luc parasite across the placental barrier, pre-treated or not with IFN-γ [100 ng/ml] or Nef [500 ng/ml]. (*n=3*, means ± SD). One-way ANOVA with Dunnett's multiple comparisons, ns (not significant), $^{**}P = 0.0072$, $^{****}P < 0.0001$. **(B)** Permeability to 40kDa FITC-Dextran post-transmigration, of RH-Luc parasite across the placental barrier, pre-treated or not with IFN-γ [100 ng/ml] or Nef [500 ng/ml]. (*n=3*, means ± SD). One-way ANOVA with Dunnett's multiple comparisons, ns (not significant), $^{*}P = 0.0143$.

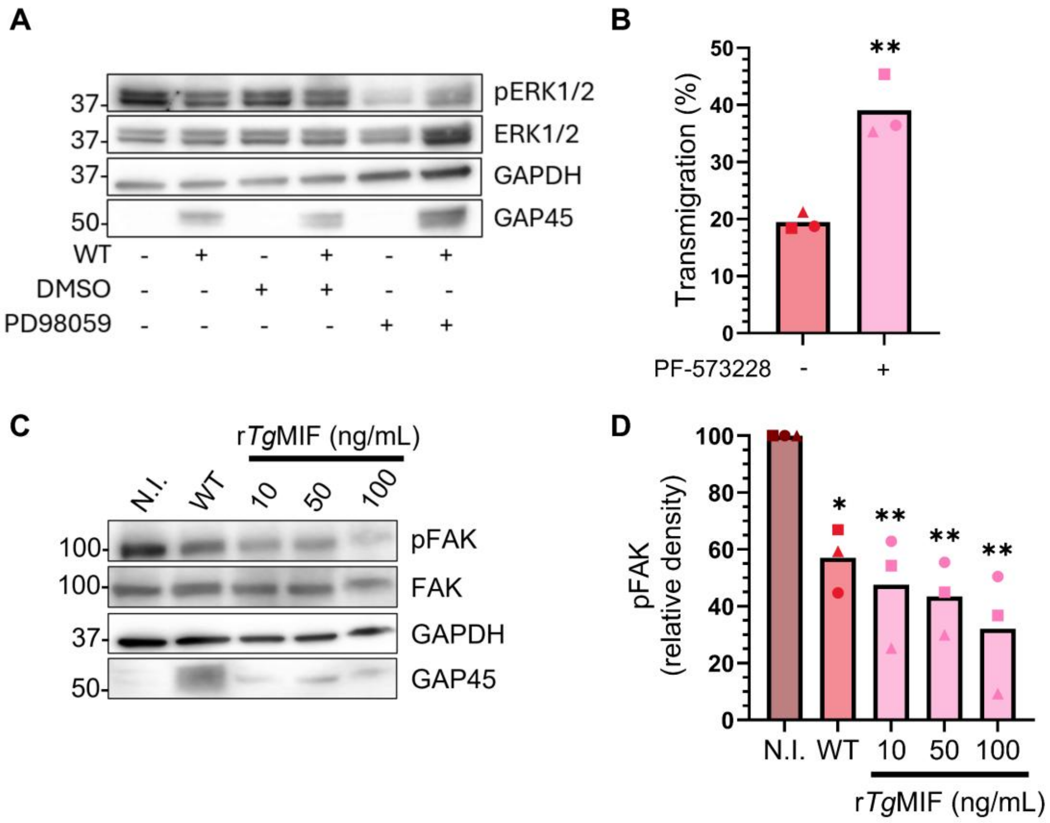


**Supplemental Figure 3. (A)** SDS-PAGE analysis of lysates from placental barriers 24 hours pre-treatment with or without PD98059 [50 µM] or DMSO vehicle, then infected or not for 45 minutes with WT (RH-Luc) parasite. The blotting was performed with anti-phospho ERK1/2 (pERK1/2), anti-ERK1/2, anti-GAPDH (cell loading control), and anti-GAP45 (parasite loading control), as well as their respective secondary HRP-antibodies. **(B)** WT parasite transmigration across placental barriers 16 hours pretreated with PF-573228 [12.5 µM]. (*n=3*, means ± SD), student's t-test, $^{**}P$ = 0.0040. **(C)** SDS-PAGE analysis of lysates from placental barriers 5 hours infected or not (NI) with WT parasite or treated with increased concentrations of r*Tg*MIF. The blotting was performed with anti-phopspho FAK (pFAK), anti-FAK, anti-GAPDH (cell loading control), and anti-GAP45 (parasite loading control), along with their respective secondary HRP antibodies. **(D)** Relative quantification of pFAK in each condition from (C) after normalization with GAPDH. (*n=3*, means ± SD). One-way ANOVA with Dunnett's multiple comparisons, ns (not significant), $^{*}P$ = 0.0280, $^{**}P < 0.01$.

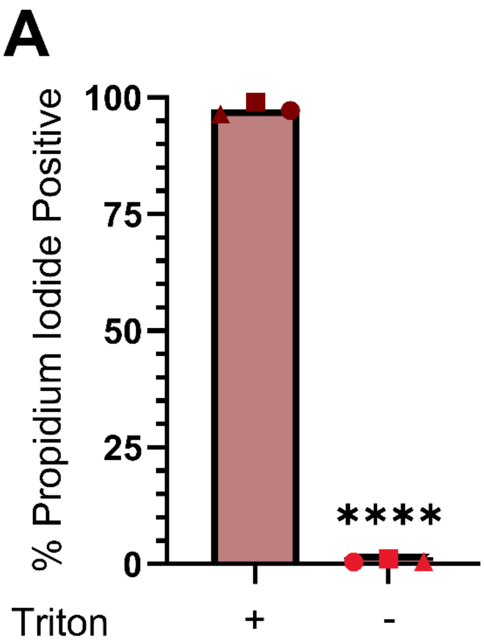


**Supplemental Figure 5. (A)** The graph represents extracellular parasite survival during excretion assay measured by propidium iodide staining, with triton-treated parasites as a death control. (*n=3*, means ± SD). student's t-test, ****$P < 0.0001$.
